# Distribution Pattern and Diversity Maintenance Mechanism of the Denitrifier Community in Alkaline Copper Mine Drainage

**DOI:** 10.1101/531434

**Authors:** Jinxian Liu, Cui Li, Zhengming Luo, Xue Wang, Jiabing Bao, Baofeng Chai

## Abstract

Denitrifiers have been shown to vary in abundance and diversity across the environmental gradients; however, the factors that determine the type and magnitude of the response of the denitrifier communities remain unclear. In order to reveal the effects of alkaline mine drainage (AlkMD) on the denitrifier community composition and diversity maintenance mechanism in a tailings reservoir, we examined the diversity patterns and adaptation mechanisms of different denitrifier communities in Shibahe copper tailings water in Zhongtiaoshan, China. The results showed that the α-diversity of *Proteobacteria* and the *nosZ*_*I*_-bearing denitrifier community in downstream seeping water (SDSW) were the highest, while those in the upstream tailings water (STW1) were the lowest. The highest *nirS* and *nirK*-bearing denitrifier community α-diversity were measured in STW1, and the lowest in STW3. The community dissimilarity of the denitrifier communities was strongly correlated with environment parameters, but the factors varied in different groups. The *Proteobacteria* community was correlated strongly with nitrate (NO_3_^-^); the *nirS* and *nirK*-bearing denitrifier communities were correlated strongly with pH, total carbon (TC), NO_3_^-^ and ammonia nitrogen (NH_4_^+^), and the *nosZ*_*I*_-bearing denitrifier community was correlated strongly with total organic carbon (TOC), dissolved oxygen (DO), NO_3_^-^, electroconductibility (EC) and spatial distance (PCNM1). Our results indicated that environmental changes were valuable index reflecting variations in denitrifier community composition in AlkMD. Additionally, the findings in this study are helpful in understanding the relationship between the microbial community composition changes and functional adaptation in different ecosystems.

**IMPORTANCE:** Metal mine tailings pond is an important site polluted by heavy metals and nitrogen, and excess nitrogen is mainly removed by microorganisms through denitrification. Although many studies have reported that *Proteobacteria* is the main denitrifying bacteria, and *nirS, nirK and nosZ-bearing denitrifiers* are three key functional groups in denitrification process. However, less is known of their response to discharge of alkaline copper tailings wastewater (AlkMD). AlkMD exerts an adverse impact on composition and structure of denitrifying bacteria community. The change of nitrogen and carbon concentration along the flow direction of wastewater resulted in the change of denitrifying bacteria community diversity pattern, that is to say, environmental filtering force drove the denitrifier community structure in this highly polluted ecosystem.

## INTRODUCTION

Understanding and inferring the factors that determine the composition and structure of microbial communities is a critical research question and an important step in understanding how functional microorganisms contribute to ecosystem processes (1). In addition to a variety of heavy metals, copper mine drainage contains a variety of inorganic pollutants such as chlorides, sulfides and nitrides. Nitrides in mine drainage are mainly derived from using explosives (2) and nitrogen-bearing flotation agents in the mining process. Nitrogen is an important contaminant in wastewater, and the toxicity of ammonia can cause aquatic animals to die. Microbial-led denitrification is the most important route for excess nitrogen removal in natural ecosystems (3), for example, ocean (4), lakes (5, 6), wetlands (7, 8) and wastewater (9, 10). In the process of denitrification, nitrates (NO_3_^-^) and nitrites (NO_2_^-^) are reduced to gaseous end products: nitric oxide (NO), nitrous oxide (N_2_O) and nitrogen (N_2_) (11). If the denitrification process is incomplete, then NO and/or N_2_O will be produced and released into the atmosphere; in particular, the powerful greenhouse gas N_2_O is the dominant ozone-depleting agent that is 310 times more potent than carbon dioxide (CO_2_) (12, 13). The activity of nitrous oxide reductase encoded by the *nosZ* gene determines whether there is excessive N_2_O emission. It has been shown that the abundance of the denitrification function genes, such as *nirS, nirK* and *nosZ* was related to the potential rate of denitrification (6, 14), so the community structure and diversity pattern of *nirS, nirK* and *nosZ*-bearing denitrifiers would have an effect on the denitrification process. Denitrifiers are environment-specific (6), and the composition and distribution of denitrifiers are significantly affected by dissolved oxygen, pH, organic carbon and heavy metals (15-18). Our previous research (19) showed that there was a significant nitrogen pollution gradient in alkaline mine drainage (AlkMD), and the copy number of denitrification genes exhibited obvious changes along the environmental gradient. However, whether the structure of the denitrifier community also changes significantly along an environmental gradient remains unclear.

Bacteria are the most important denitrifying microorganisms in many ecosystems (10, 12) and most of the known denitrifying functional gene sequences (*nirS, nirK* and *nosZ*) are derived from bacteria of the phylum *Proteobacteria* (6, 10, 20). Moreover, the composition and diversity of the *Proteobacteria* community vary significantly in various industrial wastewaters, depending on the mine drainage characteristics, treatment configurations and operating conditions (10). Linking the functional genes community composition and major denitrifying bacterial community composition would be better understand the denitrification processes controled by microbial. Some studies have shown that the structural change of denitrifying bacteria community is caused by environmental change (6, 9, 21, 22). In fact, it is still unclear, whether the functional genes community composition and major denitrifying bacterial community composition have the same adaptability to environmental changes in AlkMD.

A change in the microbial community composition and structure is an external manifestation of adaptation to the environment. Functional gene composition have been used to better understand the microbial control of biochemical processes. The structure and diversity of denitrifier communities determine the removal of excess nitrogen in ecosystems and the resilience of the polluted environment. This study aimed to address the following questions: (1) How does AlkMD affect the community structure of the major denitrifying bacteria (*Proteobacteria*)? (2) How does AlkMD affect the *nirS, nirK* and *nosZ*_*I*_-bearing denitrifier community structure? (3) How can the diversity pattern of denitrifier communities be maintained in an AlkMD environment?

## RESULTS

### Composition and structure of the Proteobacteria

*Proteobacteria* was the most abundant (the mean relative abundance accounted for 58.95% of) group in the entire bacterial community (Fig. 1), indicating that it may play an important role in maintaining the stability of the community structure. A total of 24538, 14285, 14586, 10651 and 8231 filtered sequences were obtained from the five respective samples. According to sequence alignment at a sequence identity cut-off value of 97%, 237, 406, 514, 418 and 515 corresponding OTUs were identified (Table S2). Sequence number gradually decreased along the water flow direction; however, the OTU numbers were increased.

**FIG 1.**
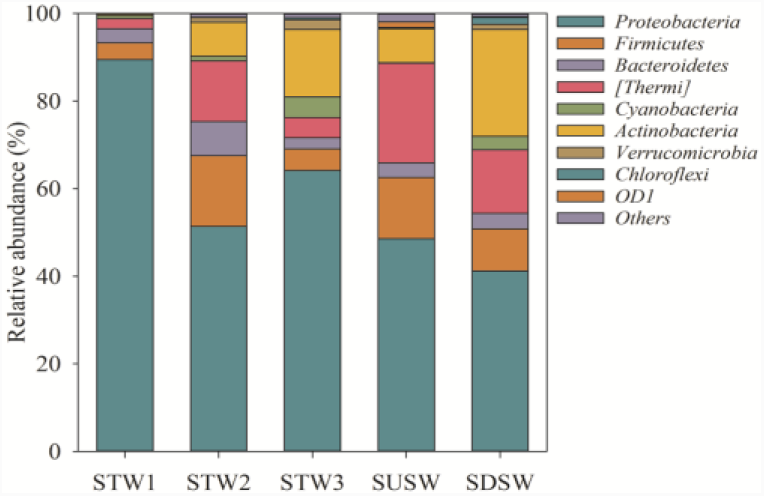
Dominant phylum composition of bacterial community in five sampling points.

The diversity index (OTUs, Chao-1, Simpson and Shannon) of the *Proteobacteria* increased gradually along the direction of the water flow (Table S3). In the three tailings reservoir water samples (STW1, STW2 and STW3), the Shannon and Simpson values were the highest in STW2. There were also differences in SUSW and SDSW, and the α**-**diversity (except sequence number) in SDSW was higher than that in SUSW (Table S2). The α**-**diversity indices of *Proteobacteria* were significantly correlated with environmental parameters (TC, pH, NO_3_^-^, NO_2_^-^ and SO_4_^2-^) (Fig. S1), indicating that the environmental gradient had a significant influence on the community α**-**diversity.

According to the literatures (listed in supplementary materials), we have screened out the main denitrifying bacterial groups in *Proteobacteria* from the class to the genus level (Fig. 2). There were mainly 4 classes (*α,β,γ,δ*-*Proteobacteria*) of *Proteobacteria* in the five samples. *α-Proteobacteria, β-Proteobacteria* and *γ-Proteobacteria* were distributed across the five samples; however, *δ-Proteobacteria* were detected in four samples except STW1 (Fig. 2a). At the order level, 11 orders are common denitrifying bacteria groups, and their average relative abundance accounts for 49.5% of the entire bacterial community (Fig. 2b). At the family levels, the average relative abundance of 19 denitrifying bacteria groups accounts for 44.91% of the entire bacterial community (Fig. 2c). At the genus level, the average relative abundance of 27 denitrifying bacteria groups accounts for 31.83% of the entire bacterial community (Fig. 2d). From order level to genus level, the relative abundance of dominant groups in *Proteobacteria* changed obviously. The relative abundance of these groups had a significant correlation with the environmental parameters (especially pH, NO_3_^-^, NO_2_^-^, TC and As) (Table S4–S6). The results of PERMANOVA showed that the *Proteobacteria* community structures at different sampling locations were different (Table S8).

**FIG 2.**
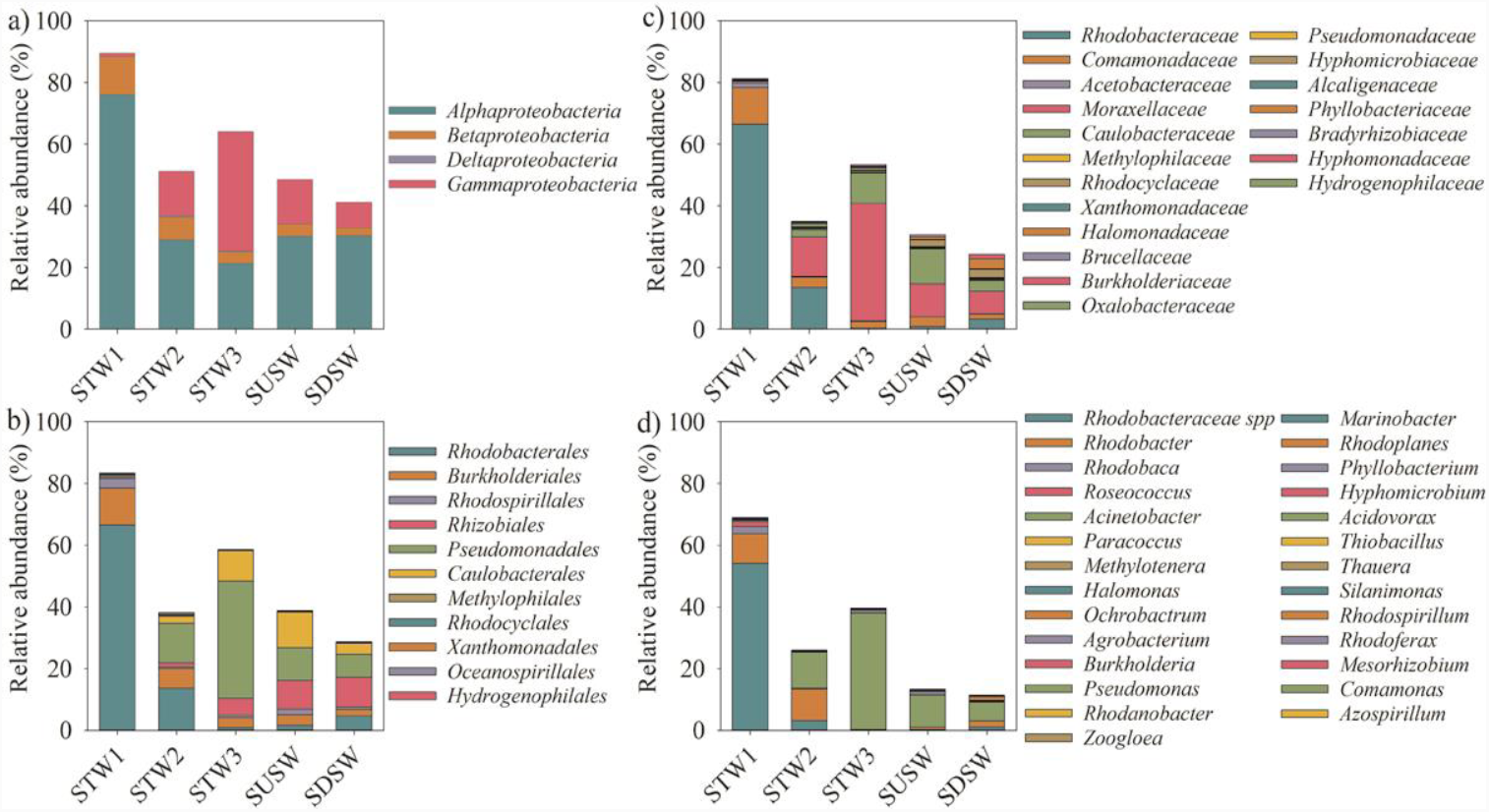
The Community composition of denitrifying bacteria in *Proteobacteria*. a) class level, b) order level, c) family level d) genus level.

### Composition and structure of nirS, nirK and nosZ_I_-bearing denitrifier communities

The composition and diversity of the *nirS, nirK* and *nosZ*_*I*_-bearing denitrifier communities showed significant differences among the five sampling points (Fig. 3, Table S7 and Table S8). The *nirS*-bearing denitrifier community a total of 33 bands (OTUs) were detected, of which 15 OTUs were dominant groups (relative abundance of more than 1%) (Fig. 3a); a total of 44 OTUs were detected in the *nirK*-bearing denitrifier community, of which 25 OTUs were dominant groups (Fig. 3b); a total of 31 OTUs were detected in the *nosZ*_*I*_-bearing denitrifier community, of which 15 OTUs were dominant groups (Fig. 3c). The mean relative abundance of dominant groups of the *nirS, nirK* and *nosZ*_*I*_-bearing denitrifier was 85.09%, 85.49% and 88.84%, respectively (Fig. 3).

**FIG 3.**
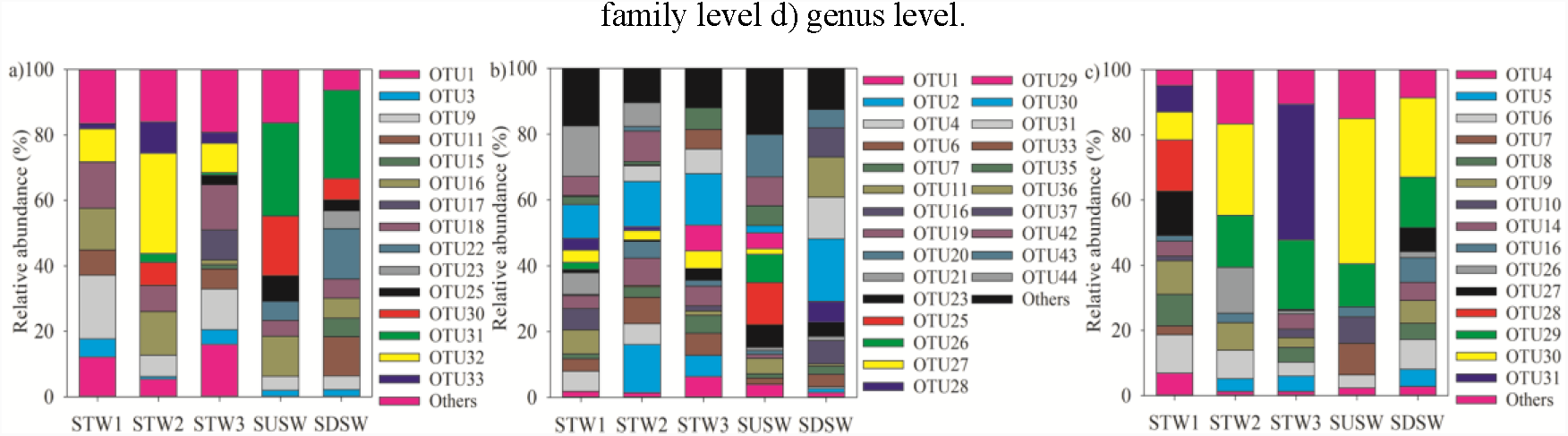
Community composition of dominant denitrifier.a) *nirS-*denitrifier community, b) *nirK-*denitrifier community, and c) *nosZ*_*I*_*-*denitrifier community.

The *nirS* and *nirK*-bearing denitrifier communities had similar α-diversity variation tendency, and their abundance gradually increased along the direction of water flow, but the richness and diversity gradually decreased. However, the abundance, richness and diversity of *nosZ*_*I*_-bearing denitrifier community were all gradually increased along the direction of water flow (Table S7). The α-diversity indexes of three denitrifier communities in SDSW were higher than those in SUSW (Table S7). There was a significant correlation between community α-diversity and environmental factors (Fig. S2). The TOC content had a significant correlation with the α-diversity indexes of the *nirS*-bearing denitrifier community (Fig. S2a). The α-diversity indexes of the *nirK*-bearing denitrifier community were significantly correlated with NH_4_^+^, SO_4_^2-^, Zn, Pb, EC and PCNM1 (Fig. S2b), and the *nosZ*_*I*_-bearing denitrifier community α-diversity indexes were significantly correlated with TC, IC, NH_4_^+^, Zn, EC and PCNM1 (Fig. S2c).

### The adaptation mechanisms and interrelationships of different communities

RDA results showed that the community structure of *Proteobacteria* was significantly affected only by NO_3_^-^, and the adjusted explanatory rates was 23.0% (F=2.2, *P*<0.05) (Fig. 4a). There were five environmental factors (pH, As, IC, DO and NH_0_^+^) had significant effects on the community structure of *nirS*-bearing denitrifierand, and the adjusted explanatory rates was 70.6% (F=7.7, *P*<0.01) (Fig. 4b), among which pH had the greatest impact (explains 41%) (Table S9). A total of four environmental factors (NO_2_^-^, PCNM1, pH and TOC) had a significant impact on the community structure of *nirK*-bearing denitrifier, and the adjusted explanatory rates was 52.8% (F=4.9, *P*<0.01) (Fig. 4c), of which NO_2_^-^ had the greatest impact (explains 23.5%) (Table S9). *nosZ*_*I*_-bearing denitrifier community structure was mainly influenced by TOC, DO, NO_3_^-^, IC, EC and PCNM1, and the adjusted explanatory rates was 75.6% (F=8.2, *P*<0.01) (Fig. 4d), of which TOC had the greatest impact (explains 25.7%) (Table S9).

**FIG 4.**
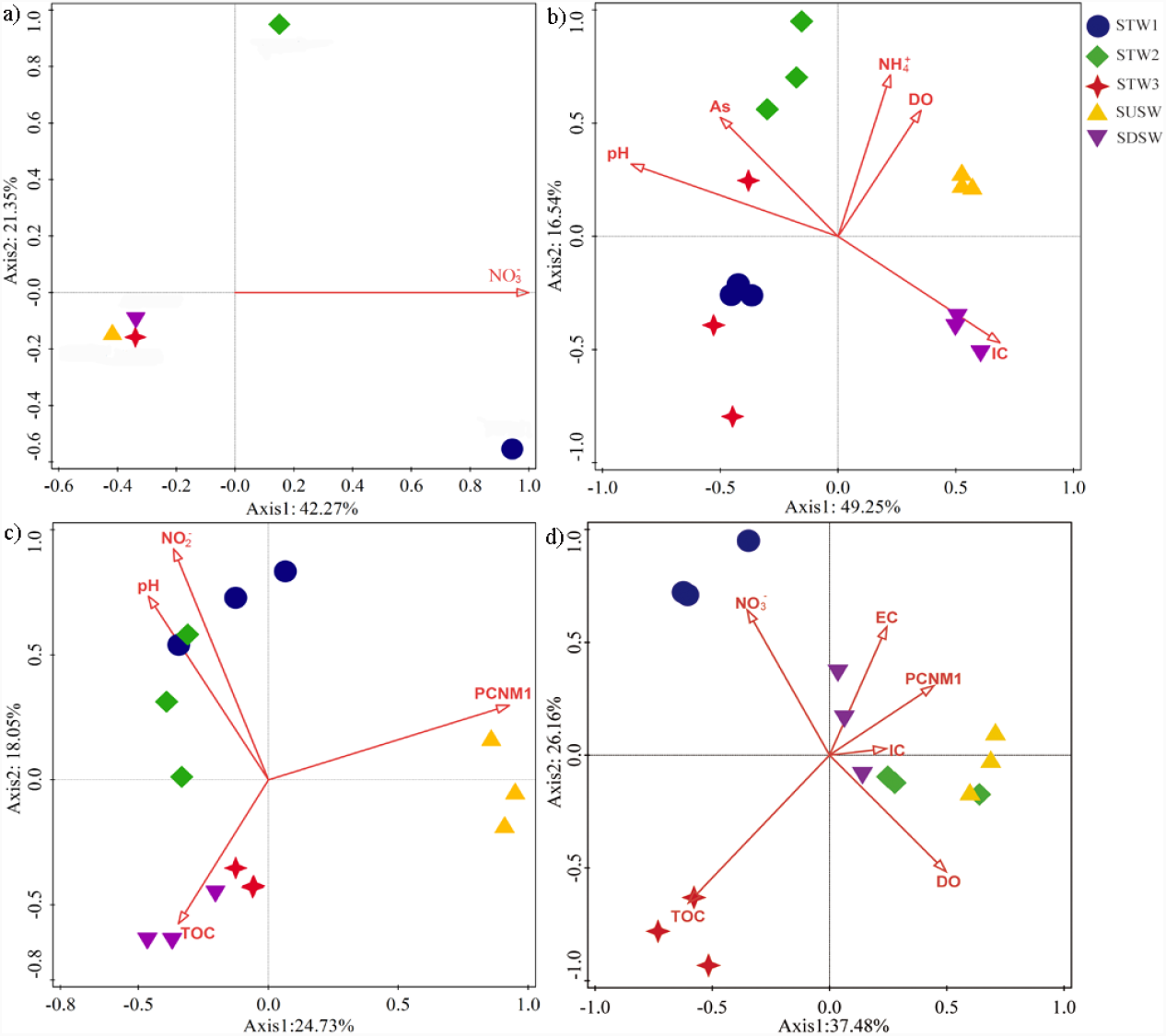
Redundancy analysis based on Bray-Curtis similarities of bacterial communities and Euclidean-Metric of environmental parameters in five sampling points. a) *Proteobacteria* community, b) *nirS-*denitrifier community, c) *nirK-*denitrifier community, and d) *nosZ*_*I*_*-*denitrifier community.

**FIG 5.**
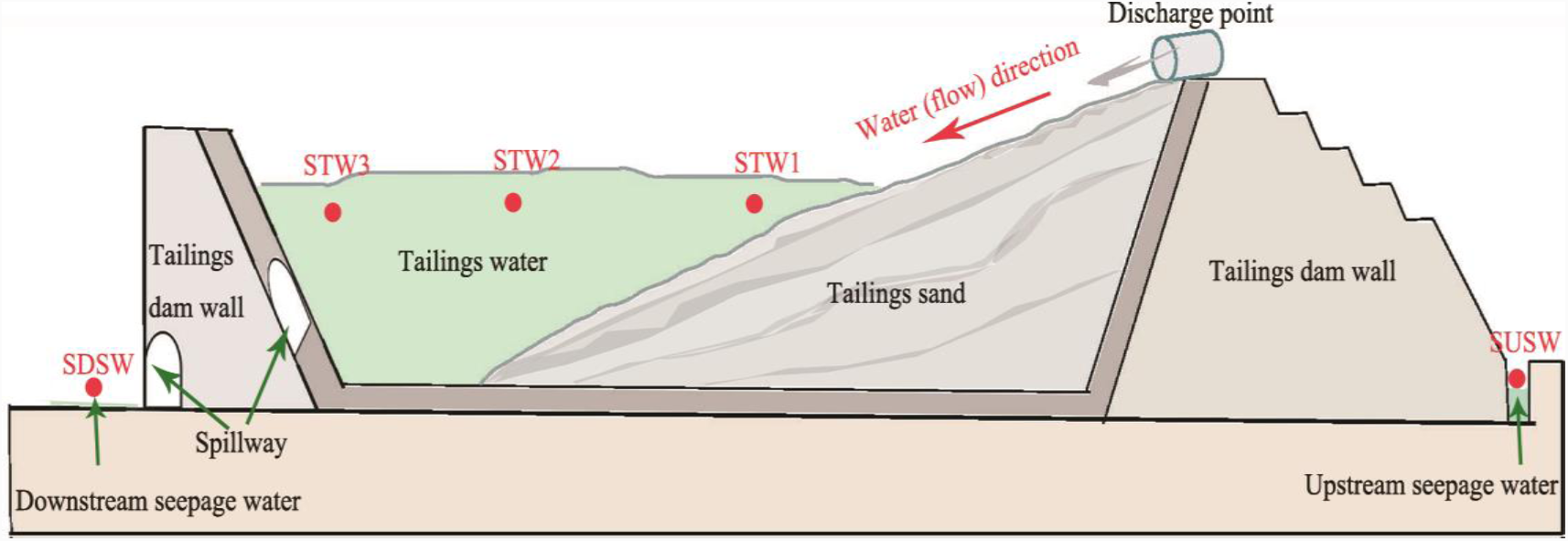
Sketch map of sampling points for Shibahe tailings reservoir. STW1, STW2 and STW3 are three tailings reservoir water sampling point; SUSW and SDSW are upstream and downstream seepage water sampling points, respectively.

The community structures of *nirS* and *nirK*-bearing denitrifier community were significantly correlated with *Proteobacteria* community structure, however the community structure of *nosZ*_*I*_*-*bearing denitrifier was significantly correlated with entire bacterial community structure (Table S10). The community structure of *nirS*-bearing denitrifier was significantly correlated with *β* and γ-*Proteobacteria* community structure, and the community structure of *nirK*-bearing denitrifier was significantly correlated with *α* and γ-*Proteobacteria* community structure, however the community structure of *nosZ*_*I*_*-*bearing denitrifier was only significantly correlated with γ-*Proteobacteria* community structure (Table S11).

Analysis of variation partitioning (VPA) between environmental factors and spatial distance was used to examine their relative influence on the community structure of the three enitrifier communities and *Proteobacteria*. Environmental factors and spatial distance together explained 61.5%, 39.9%, 71.6% and 33.7% of the *nirS, nirK, nosZ*_*I*_-bearing denitrifier communities and *Proteobacteria* respectively (Table 1). Environmental factors had a significant influence on the four communities and independently explained 55.5%, 26.6%, 60.5% and 27.8% respectively, while independent spatial distance had a significant influence on the structure of *nirK* and *nosZ*_*I*_-bearing denitrifier community (Table 1). The interaction effect between spatial distance and environmental factors also had an effect on community assembly except for *nirK*-bearing denitrifier community (Table 1).

**TABLE 1.**
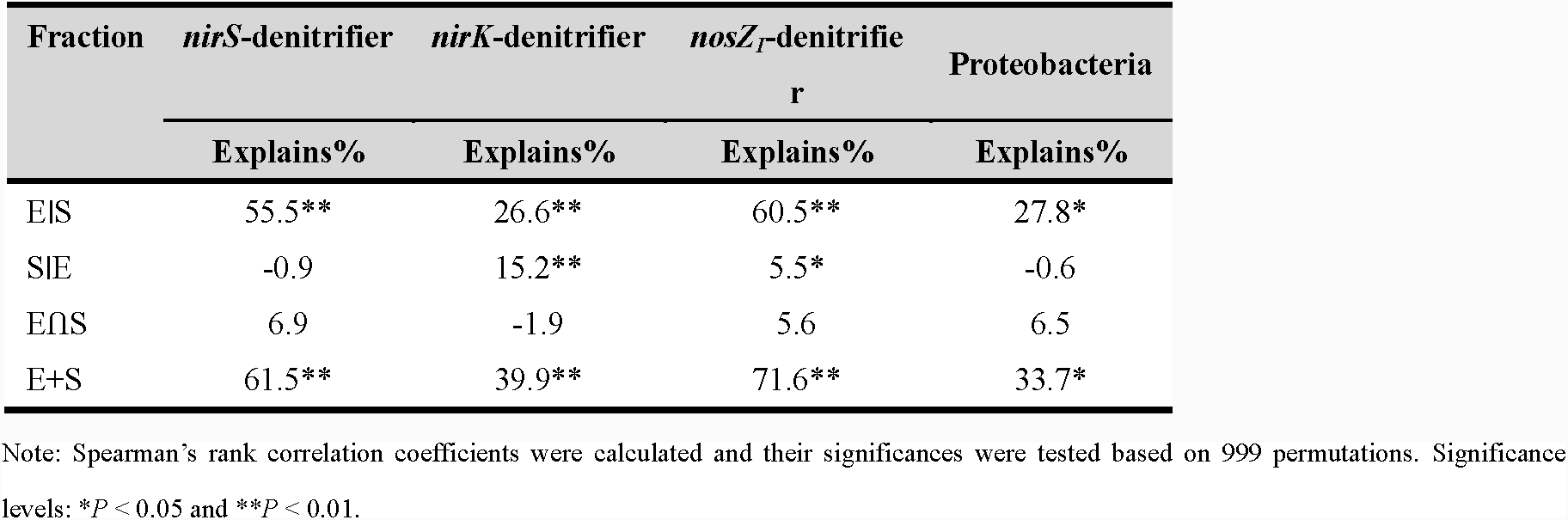
Variation partitioning results showing the contribution of the exclusive environmental (E|S), exclusive spatial (S|E) factors, as well as the interaction effect components (E⋂S) and the together effect (E+S) of the four communities based on the Bray-Curtis distance.

## DISCUSSION

### Composition and diversity of the Proteobacteria community in AlkMD

*Proteobacteria* is the major denitrifying bacterial groups (6, 10) that also are important components of the bacterial community in this study (average abundances were 58.95%) (Fig 1). Some studies showed that *Proteobacteria* is sensitive to changes in local environmental conditions (6, 19, 23), and our results were consistent with these previous studies. Our data showed that the diversity of denitrifying bacteria belonging to the *Proteobacteria* (from class to genus) changed along the flow direction of drainage (Fig. 2). The differences in environmental adaptability, such as responses to the availability of nitrate, sulfate and carbon for energy metabolism, and the resistance to an AlkMD environment, leads to different composition patterns of *Proteobacteria* community in five samples. For example, *Rhodobacteraceae* and *Rhodobacte* are photoautotrophic bacteria, and nitrate is an electron acceptor during anaerobic respiration (10); *Legionella* is sensitive to changes in dissolved organic carbon concentrations (24), while, *Hydrogenophaga* can better adapt to the environmental contaminated by nitrogen and sulfur (25).

Not all denitrifying bacteria groups were significantly correlated with NO_3_^-^ or NO_2_^-^ concentration (Table S4-S6), suggesting that different denitrifying bacteria play different roles in denitrification process (6, 26). There was a significant positive correlation between the relative abundance of *Oceanospirillales* and NH_4_^+^ concentration (Table S4), because they obtain energy via conversion of NH_4_^+^ to NO_2_^-^ (27). Of course, pH and carbon source were also important factors affecting the composition and structure of denitrifying bacteria community (Fig. S1and TableS4-S6) (10, 15, 28). Extreme pH levels (>8.5 or <4) cause changes in microbial cell membrane permeability and eventually leads to apoptosis; while carbon source is the main energy source of denitrifying bacteria, and the denitrification rate will be significantly reduced in environment with insufficient carbon source(10). Many studies have reported that microorganisms were sensitive to heavy metal stress, especially in industrial wastewater (19, 23, 29, 30). Although heavy metals may change the composition and spatial structure of bacterial communities, it is interesting to note that the abundance of *Proteobacteria* increased as the As content increased (Fig. S1). This phenomenon was discovered in other studies (31, 32). This may be explained by that the higher pH (pH>8) in this study area, heavy metal ions form insoluble compounds, reducing bioavailability (31).

RDA results showed that NO_3_^-^ concentration was the main and only factor affecting the spatial distribution diversity of *Proteobacteria* community (Fig. 4a, Table S9). Most of denitrifier communities are taxonomically affiliated with *Proteobacteria* (10) and Yoshida (11, 22) showed that *Rhodobacteraceae, Rhodobacter, Burkholderia* and *Roseococcus* are denitrifying bacteria containing *nirS* or/and *nirK* functional genes. In this study, these nitrite-reducing bacteria are dominant genera in *Proteobacteria* (Fig. 2d), and the diversity pattern of dominant groups often determines the distribution pattern of the whole community. Therefore, it is not difficult to understand that the change of nitrate concentration was the main reason affecting the diversity of the spatial distribution pattern of *Proteobacteria* community.

### Composition and diversity of nirS, nirK and nosZ_I_-bearing denitrifier communities in AlkMD

*nirS, nirK* and *nosZ*_*I*_-bearing denitrifiers are particularly important in the denitrification pathway because *nirS* and *nirK* genes code for nitrite reductase that catalyzes the first committed step to a gaseous product (NO_2_^-^ → NO) (33), and greenhouse gas nitrous oxide reduction (N_2_O → N2) catalyzed by nitrite reductase encoded by *nosZ*. *nosZ* genes actually belong to two phylogenetically distinct clades (*nosZ*_*I*_ and *nosZ*_*II*_) (34, 35); unfortunately, *nosZ*_*I*_ was successfully amplified but *nosZ*_*II*_ amplification failed in this study. The composition, structure and diversity of *nirS, nirK* and *nosZ*_I_-bearing denitrifier communities differed at five sampling points (Fig. 3 and Tables S7, S8), and this diversity pattern was significantly influenced by the changes of environmental factors (Fig. 4 and Fig. S2). Jones (36) found that the *nirK* gene was contained in a broader range of taxa, and nearly one-third of denitrifiers do not contain the *nosZ* gene (37). This may be the main reason for the lower diversity of *nosZ*_I_-bearing denitrifier community. The diversity change trends of the *nirS* and *nirK*-bearing denitrifier communities were similar (Table S7), because these two groups have the same ecological function in the denitrification process (6, 12). The community structures of *nirS, nirK* and *nosZ*_*I*_-bearing denitrifier were believed to be controlled by different environmental factors (Fig. 4b,c,d). This was similar with some previous studies (6, 38), NH_4_^+^ had a significant effect on the structure of *nirS*-bearing denitrifier community (Fig. 4b), possibly due to the most abundant *nirS* sequences were matched *β-Proteobacteria* and *γ-Proteobacteriain* (Table S11) (6). Because *β* and *γ-Proteobacteria* are also the main ammonia-oxidizing bacteria, they oxidize ammonium to nitrite to obtain energy (27). Dissolved oxygen also had a significant effect on the community structure of *nirS* and *nosZ*_*I*_-bearing denitrifier (Figs. 4b and 4d). Our results also showed that *nirS*-bearing denitrifier community had higher diversity in STW1 with lower DO, but *nosZ*_*I*_-bearing denitrifier community had higher diversity in SDSW with higher DO (Table S1, Table S7). Some studies have shown that most of *nirS*-bearing denitrifier are anaerobic or facultative anaerobes, while *nosZ*_*I*_-bearing denitrifier are aerobe (14, 17, 18). The effect of pH, on *nirS* and *nirK*-bearing denitrifier community structure was also significant (Figs. 4b and 4c). The optimum pH range of denitrifying bacteria is 6.0-8.5, beyond which the structure and function of denitrifying bacteria will be significantly affected (39). In this study, the range of pH was 8.01-9.38(Table S1), so the denitrifier community structure were significantly affected. The substrates NO_2_^-^ and/or NO_3_^-^ are used as electron acceptors in denitrification, and TOC is the main energy source and electron donor of denitrifier, so these two factors have a significant impact on the community structure of *nirK* and *nosZ*_*I*_-bearing denitrifier (6, 40). While, the energy source of *nirS*-bearing denitrifier may be mainly come from ammonia oxidation process (27, 41). Interestingly, spatial distances also had a significant impact on the *nirK* and *nosZ*_*I*_-bearing denitrifier community structure (Table 1), indicating that in addition to environmental choices, dispersal limitation was also an important reason of differences in spatial distribution of communities.

### Community correlation and diversity maintenance mechanism of denitrifiers

The community structure of *nirS* and *nirK*-bearing denitrifiers was significantly correlated with *Proteobacteria* community structure, and *nosZ*_*I*_-bearing denitrifiers community structure was significantly correlated with entire bacterial community structure (Table S10). The results of Yoshida’s (11, 22) studies were similar to our’s study results, suggesting that most *nirS* and *nirK*-bearing denitrifiers are *Proteobacteria* in both paddy soil and tailings wastewater. While, *nosZ* gene fragments were found in *Proteobacteria, Bacteroides, Firmicutes* and other bacterial groups (*Chloroflexi* and *Actinobacteria*) (6, 42-44), so *nosZ*_*I*_-bearing denitrifiers community structure changed with the change of the entire bacterial community structure. The community structure of *nirS, nirK* and *nosZ*_*I*_-bearing denitrifiers were all significant correlation with the community structure of *γ-Proteobacteria* (Table S11), indicating that *γ-Proteobacteria* play an important role in the whole denitrification process in this study. Whether*γ-Proteobacteria* is a complete denitrifiers remains to be further studied. Although the *nirS* and *nirK* gene fragments encode functionally identical enzymes, *nirS* and *nirK*-bearing denitrifiers are composed of different microorganisms. The community structure of *β-Proteobacteria* was significantly correlated with the community structure of *nirS*-bearing denitrifiers, while the community structure of *α-Proteobacteria* was significantly correlated with the community structure of *nirK*-bearing denitrifiers (Table S11). This result is consistent with Liu’s research results on denitrifying bacteria communities in the sediments of freshwater lakes. Saarenheimo’s (6) research also showed that *nirS* sequences matched *β-Proteobacteria* and the most abundant *nirK* sequences matched *α-Proteobacteria*.

The mechanism underlying the denitrifier community composition and the associated processes that respond to environmental gradients is the core of research in nitrogen-contaminated habitats. In this study, we found that environmental selection drove the diversity pattern of the *nirS, nirK* and *nosZ*_*I*_-bearing denitrifiers and *Proteobacteria* communities (Table 1). Although dispersal limitation also had an effect on the spatial pattern of the *nirK* and *nosZ*_I_-bearing denitrifier community, the intensity of environmental selection was much greater (Table 1).

### Conclusion

In this study, we observed that denitrifier community composition and spatial diversity pattern were mainly dependent on the environmental conditions, which changed along the water flow through the tailings reservoir, further implying environmental selection was the main reason for maintaining the diversity pattern of the denitrifier community in AlkMD. The diversity of the denitrifier communities was strongly controlled by the carbon, nitrogen and pH, suggesting that the importance of niche-based processes in microbial community assembly may strongly depend on the local environmental context. Our results thus highlighted that an environmental filtering force drove the denitrifier community structure in this highly polluted ecosystem. To further reveal how environmental changes affect the diversity of denitrifier communities, the next step is to better understand how the metabolic activities of denitrifier communities change in AlkMD.

## MATERIALS AND METHODS

### Site description and sampling procedures

Shibahe tailings reservoir (35°15′∼35°17′N, 111°38′∼111°39′E) is a valley-type tailings reservoir surrounded by tailings dam, flood dam and mountain body on both sides. All The length of tailings reservoir is about 3600 m, the length of catchment is about 800 m, and the average width is 600 m. In the middle zone line of catchment area, three tailings reservoir water sampling points (STW1, STW2 and STW3) were selected along the flow direction, the interval of sampling points was 350 meters, and also two seeping water sampling points (SUSW and SDSW) were selected (Fig.5). Detailed sampling procedures were described in supplementary materials.

### Physiochemical analysis

Referring to methods of Liu (19), water physical-chemical parameters include pH, dissolved oxygen (DO), conductivity (EC), nitrate (NO_3_^-^), ammonium (NH_4_^+^), total carbon (TC), total organic carbon (TOC), inorganic carbon (IC), nitrite (NO_3_^-^), sulfate (SO_4_^2-^) and heavy metals (As, Cd, Cu, Pb, Zn) content were measured at each sampling point. The results of physical-chemical parameters in supplementary materials (Table S1).

### DNA extractions, PCR amplification and high-throughput sequencing

DNA of microorganisms on filter membranes were extracted using a Fast DNA SPIN extraction kits (MP Biomedicals, Santa Ana, CA, USA), then the V3-V4 hyper variable region of 16S rDNA gene and denitrification functional genes (*nirS, nirK* and *nosZ*_*I*_) of bacteria were amplified. The bacterial community structure was obtained through high throughput, while denitrifying functional community structure was obtained by denaturing gradient gel electrophoresis (DGGE). The system and conditions of PCR amplification and the analysis of high throughput data were all following the same protocol as in (19).

### PCR-DGGE analysis

For DGGE analysis, PCR amplification fragments of *nirS, nirK* and *nosZ*_*I*_ were obtained from DNA using the primer pairs listed in Table S2. A GC-clamp (CGCCGCGCGCGGCGGGCGGGGCGGGGGCACGGGGGG) was attached to the 5′ of each forward primer. Then the PCR products were analyzed by electrophoresis in 2% agarose gel, followed by DGGE which was described in detail in the supplementary materials.

### Statistical analysis

All data were normalized (square root**-** or log**-**transformed) to reduce heteroscedasticity before analysis. Statistical differences among the sampling points of environmental parameters, α-diversity of *Proteobacteria, nirS, nirK* and *nosZ*_*I*_-bearing denitrifier community were analyzed by one-way ANOVA, and the Waller-Duncan significance difference test was used for multiple comparison. Differences in the community structure of denitrifiers at different sampling points were verified by PERMANOVA. Correlations between the denitrifier community diversity (α- and β-diversity) and environmental variables were compared using Spearman’s rank correlation. The influencing factors on community structure were screened out by distance-based redundancy analysis (db-RDA), then analyze the influence of environmental factors (the results of db-RDA) on the community structure through redundancy analysis (RDA) (45). Finally, the contribution of environmental factors and spatial distance to the variation of four communities was respectively evaluated using variance partitioning analysis (VPA). Statistical analyses were carried out in SPSS 20.0 (IBM SPSS statistics, USA), CANOCO (version 5.0, USA), PAST (version 3.15) and the R environment using the vegan, SoDA, PCNM, psych, igraph (46), reldist and bipartite packages. The confidence interval of all statistical analyses were 95% (*P* < 0.05).

## ACKNOWLEDGMENTS

This study was supported by National Science Foundation of China (31772450; 31801962).

